# First high-quality genome assembly data of sago palm (*Metroxylon sagu* Rottboll)

**DOI:** 10.1101/2021.12.29.474478

**Authors:** Leonard Whye Kit Lim, Melinda Mei Lin Lau, Hung Hui Chung, Hasnain Hussain, Han Ming Gan

## Abstract

The sago palm (*Metroxylon sagu* Rottboll) is a all-rounder palm, it is both a tropical halophytic starch-producing palm as well as an ornamental plant. Recently, a genome survey was conducted on this palm using Illumina sequencing platform but the BUSCO genome completeness is very low (21.5%) and most of them (∼78%) are either fragmented or missing. Thus, in this study, the sago palm genome completeness was further improved with the utilization of the Nanopore sequencing platform that produced longer reads. A hybrid genome assembly was conducted and the outcome was a much complete sago palm genome with BUSCO completeness achieved at as high as 97.9% with only ∼2% of them either fragmented or missing. The estimated genome size of the sago palm is 509,812,790 bp in this study. A sum of 33,242 protein-coding genes were revealed from the sago palm genome and around 96.39% of them had been functionally annotated. An investigation on the carbohydrate metabolism KEGG pathways also unearthed that starch synthesis was one of the major sago palm activities. These data are indispensable for future molecular evolutionary and genome-wide association studies.

**Specifications Table:** 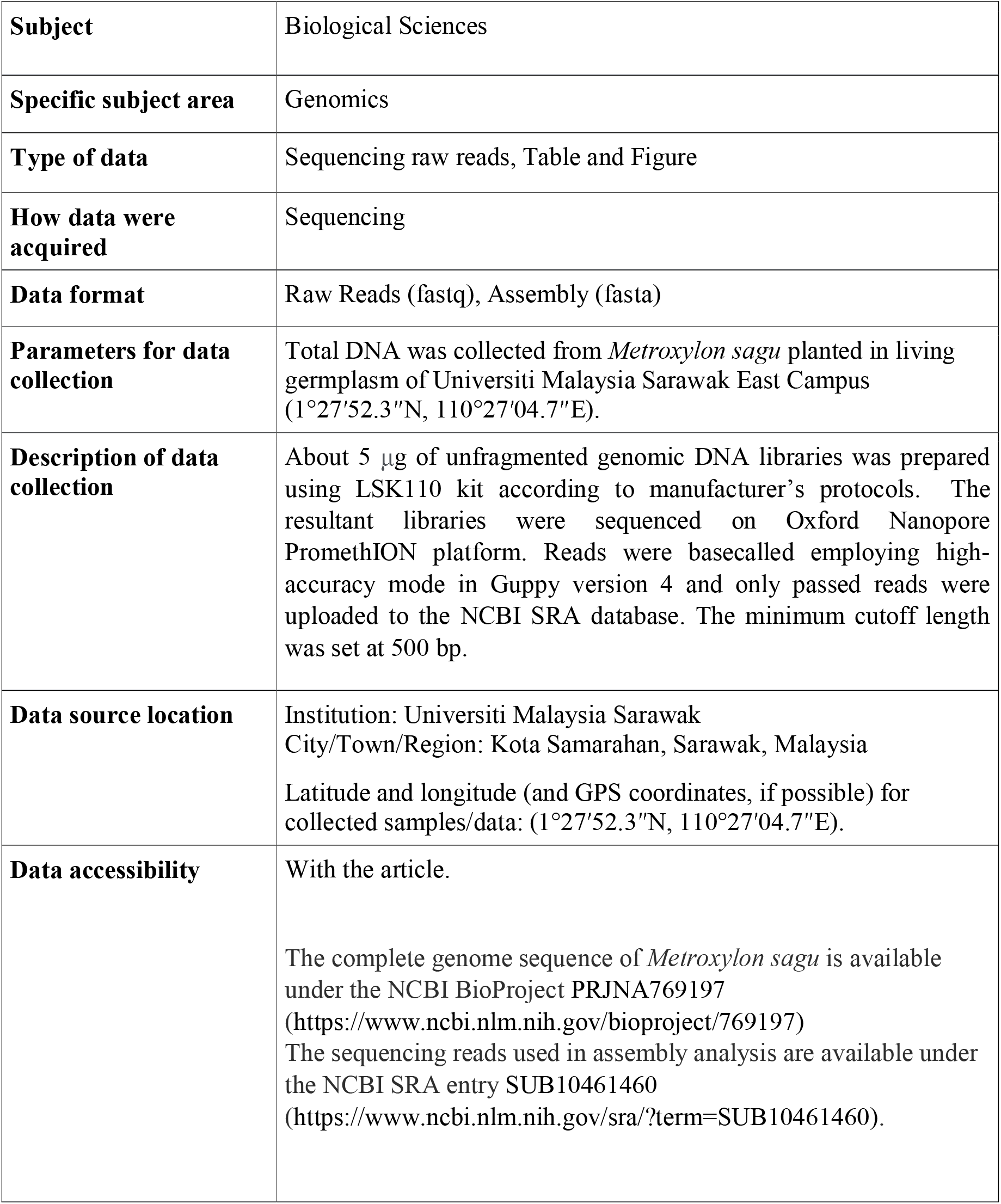

**Value of the Data:**

- First complete genome dataset for the eco-economic important sago palm (*Metroxylon sagu* Rottboll).
- High completeness of sago palm genomic dataset will facilitate future researches, such as genome-wide association studies.
- The data is useful in pioneering sago palm genetic landscape investigations which in turn unmask the mystery behind its high starch yield, salinity tolerance and disease resistance.

## Data Description

The sago palm, *Metroxylon sagu* Rottboll, is a tropical halophytic starch-producing palm well-known across communities from Papua New Guinea, Indonesia, the Philippines as well as Malaysia. This palm is an all-rounder palm, fulfilling all the required properties of a dream crop and an ornamental plant. This palm of life can be harvested from top to bottom for various imperative usage across a myriad of industries like fermentation, paper making, pharmaceutic manufacturing, biofuel production, as well as polymer and textile starting materials, to name a few (reviewed by Lim et al., 2019). The starch-producing capacity of this palm is like no other commercial food crops widely known to date as it can produce at least triple of that of rice, corn and cassava (Karim et al., 2008) besides being equipped with three times higher salt resistance than any food crops known so far (Yoneta et al., 2006; Lim & Chung, 2020).

In recent years, the sago palm has been placed in the limelight of agricultural molecular research subsequent to the whole chloroplast genome sequencing, orgnanellar genome copy number investigation, genome survey, genome size estimation, population genetics, transcriptomics as well as proteomics (Wee & Roslan, 2012; Abbas et al., 2017; Lim et al., 2020a; Lim et al., 2020b; Nisar & Hussain, 2020; Lim et al., 2021). The origin of life is often associated with the central dogma of molecular biology and genes are deemed the building blocks of life. With the next generation sequencing technologies developing at an unprecedented booming pace, it has become one of the best options to venture into the unknown genomic data of the sago palm due to its low cost and high throughput advantages. Not long ago, an interesting hybrid genome assembly approach were introduced and its efficacy in achieving higher genome completeness has been proven in species such as clown anemonefish and Murray cod (Austin et al., 2017; Tan et al., 2018). Ergo, the combinatorial power of the Illumina accurate short reads and long but less accurate Nanopore reads were emulated in this study to conduct a hybrid assembly of genome sequence of the sago palm in hope to decipher the unventured genomic landscape of this palm of life.

The sago palm genome was estimated at 509,812,790 bp in this study. The sago palm genome sequencing from this study had greatly improved the genome completeness in an unprecedented manner. Previous report on BUSCO genome completeness of sago palm by Lim et al. (2021) revealed a 21.5% single-copy complete genes and 1.1% duplicated complete genes whereas 32.2% and 45.2% are fragmented and missing respectively. In this study, the nanopore approach coupled with the Illumina method has yielded 97.9% complete genes (89.8% single-copy and 8.1% duplicated), 1.1% is fragmented as well as only 1% is missing (Table 1). The total number of contigs in this study is 2025, which is miniature to the 739,583,200 contigs reported on the sago genome survey (Lim et al., 2021). However, the overall contig lengths has significantly in this study with 1801 contigs documented to have at least 50,000 bp. On the side note, the GC content of this improved sago palm genome assembly is 36.5%, which is a little lower than that reported by Lim et al. (2021) (37.31%).

**Table 1.** Genome statistics of the sago palm (*Metroxylon sagu* Rottboll) genome.

| Organism | *Metroxylon sagu* |
| --- | --- |
| SRA | SUB10461460 |
| Bioproject | PRJNA769197 |
| Biosample | SAMN15953627 |
| Accession number | JACVBY000000000 |
| BUSCO completeness | 97.9% complete |
| (Total BUSCO groups searched: 1614) | (89.8% single-copy; 8.1% duplicated);<br>1.1% fragmented, 1% missing |
| # contigs (>= 0 bp) | 2025 |
| # contigs (>= 1000 bp) | 2025 |
| # contigs (>= 5000 bp) | 2024 |
| # contigs (>= 10000 bp) | 2022 |
| # contigs (>= 25000 bp) | 2002 |
| # contigs (>= 50000 bp) | 1801 |
| Total length (>= 0 bp) | 509812790 |
| Total length (>= 1000 bp) | 509812790 |
| Total length (>= 5000 bp) | 509810926 |
| Total length (>= 10000 bp) | 509794905 |
| Total length (>= 25000 bp) | 509389047 |
| Total length (>= 50000 bp) | 501302081 |
| # contigs | 2025 |
| Largest contig | 4418102 bp |
| Genome size | 509812790 bp |
| GC (%) | 36.50 |
| N50 | 453371 bp |
| N75 | 220193 bp |
| L50 | 341 bp |
| L75 | 742 bp |
| # N's per 100 kbp | 0.00 |
| Genome annotation |  |
| # protein-coding genes | 33,242 |
| # functionally annotated proteins | 32,041 |
| Mean protein length | 313 aa |
| Longest protein | 3768 aa<br>(titin protein) |
| Shortest protein | 23 aa<br>(TIGR01906 family membrane protein) |
| Mean exon length | 276 bp |
| Longest exon | 8579 bp<br>(heavy metal-associated isoprenylated plant protein 3) |
| Shortest exon | 2 bp<br>(uncharacterized gene) |
| Mean intron length | 926 bp |
| Longest intron | 25685 bp<br>(putative eukaryotic translation initiation factor) |
| Shortest intron | 5 bp<br>(pyruvase kinase isozyme A) |

A total of 33,242 protein-coding genes were discovered from the sago palm genome in this study. Around 96.39% (32,041) of these protein-coding genes have been successfully annotated in terms of functionality. The mean protein length reported in this study is 313 aa. The largest protein found within the sago palm genome in this study is titin protein with an amino acid length of 3768 aa whereas the smallest protein is only 23 aa, namely the TIGR01906 family membrane protein. The mean exon and intron lengths uncovered in this study are 276 bp and 926 bp correspondingly. The lengthiest exon is as long as 8579 bp, revealed within the heavy metal-associated isoprenylated plant protein 3 gene. An uncharacterized sago palm gene contains the shortest exon of 2 bp among all other exons within the genome. The longest sago palm genome intron belongs to that of the putative eukaryotic translation initiation factor gene, with 25685 bp in length. The shortest intron (5 bp) was unearthed within the pyruvase kinase isozyme A gene of the sago palm.

All protein-coding genes of sago palm were subjected to functional annotation and the focus was placed on the carbohydrate metabolism KEGG category as this is one of the most imperative yet untapped aspect of he sago palm contributing t its high starch yield. Under the carbohydrate metabolism category, there are a sum of 15 pathways described, namely glycolysis/gluconeogenesis, citrate cycle (TCA cycle), pentose phosphate pathway, pentose and glucuronate interconversions, fructose and mannose metabolism, galactose metabolism, ascorbate and aldarate metabolism, starch and sucrose metabolism, amino sugar and nucleotide sugar metabolism, pyruvate metabolism, glyoxylate and dicarboxylate metabolism, propanoate metabolism, butanoate metabolism, C5-branched dibasic acid metabolism, as well as inositol phosphate metabolism. A total of 2221 protein-coding genes were unravelled to be closely related to the 15 aforementioned carbohydrate metabolism pathways. Majority of them (260 or 12%) are related to the amino sugar and nucleotide sugar metabolism. There is equal amount of genes (256 or 11%) found to have association with the glycolysis/gluconeogenesis as well as starch and sucrose metabolism pathways. Only 1% (22) of them are involved in the C5-branched dibasic acid metabolism. This phenomenon evidenced that the starch production is the major limelight among all other carbohydrate metabolism pathways in the sago palm genome context (Nisar & Hussain, 2020).

To date, there are only 20 representative full monocot genomes with complete records of coding sequences and proteins found within the GenBank public database. The maximum likelihood phylogenetic tree was constructed based on BUSCO single-copy proteins across the full genomes of 21 monocots (including sago palm) and two dicot outgroups (Figure 2). A sum of five major clades were found within the tree plotted and each of them represent an order, namely Dioscoreales, Poales, Arecales, Zingiberales as well as Asparagales. Most clades are backed with strong bootstrap ratio of 1, indicating high bootstrap confidence. The sago palm was found grouped under the order Arecales along with two other palm members, the oil and date palm, in this study. The Arecales order was underrepresented due to the lack of available full genome sequences. More informative branching can be observed among the palm members when more full genomes are sequenced in the future as seen in the sago palm phylogenetic trees constructed based on chloroplast genome and ITS2 region (Lim et al., 2020a; Lim et al., 2021). In a nutshell, this sago palm complete genome would be a valuable resource for future molecular evolutionary studies as well as genome-wide association studies.

**Figure 1.**
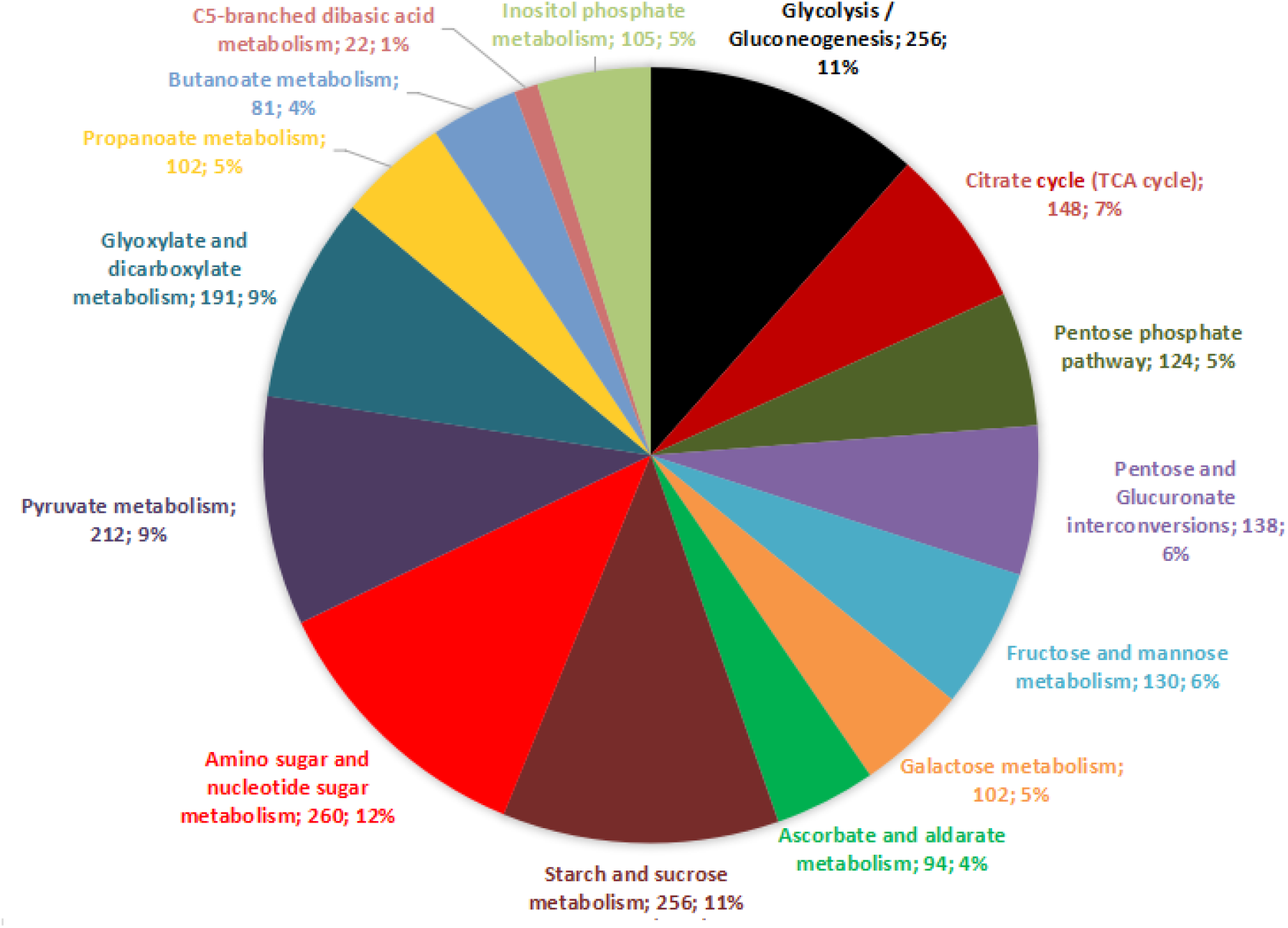
The breakdown of the KEGG carbohydrate metabolism pathways the sago palm genes are involved in.

**Figure 2.**
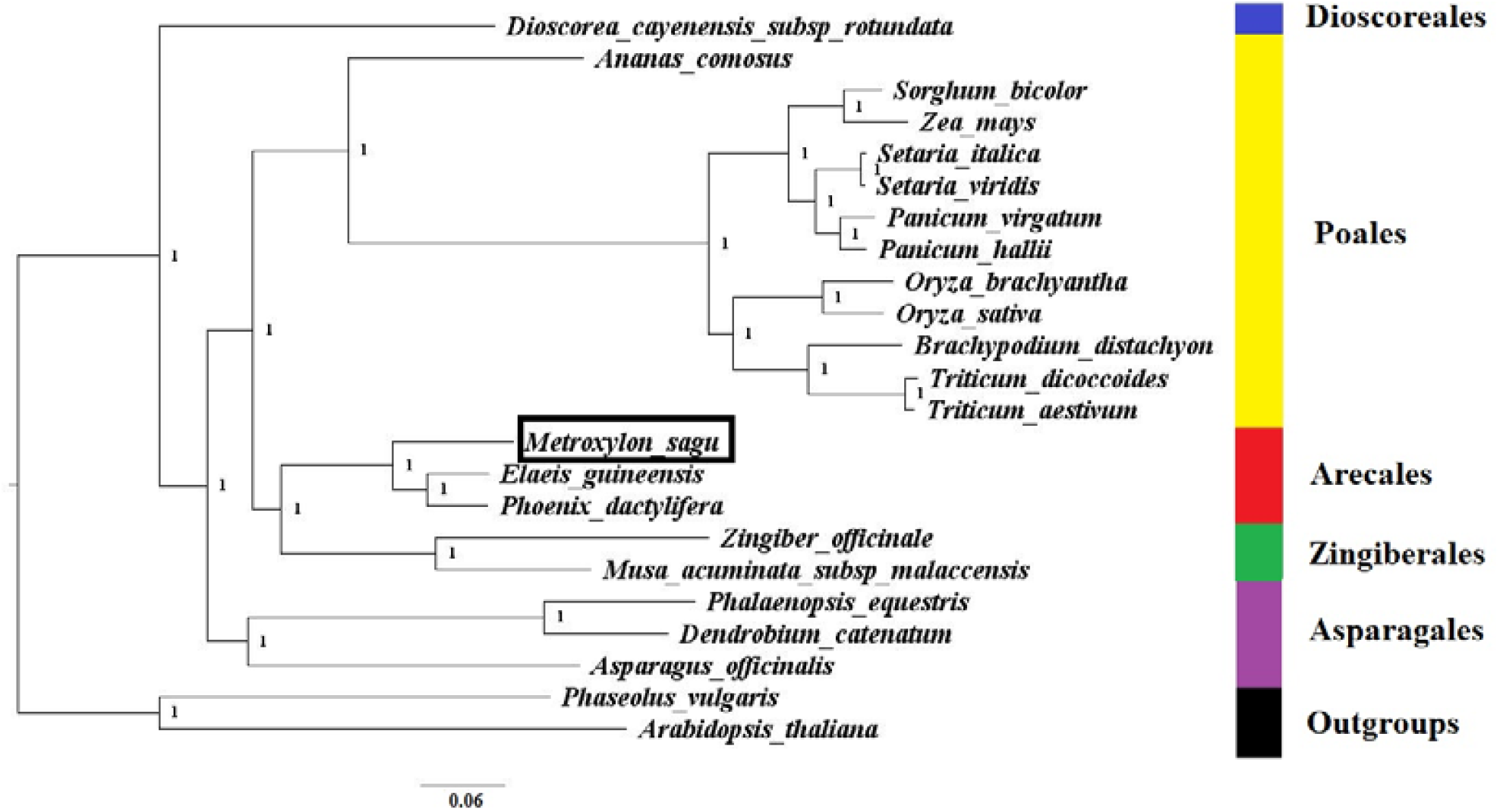
The maximum likelihood whole genome phylogenetic tree constructed based on BUSCO genes across all the available full monocot genomes (with complete records of coding sequences and proteins) found within the GenBank public database, with 1000 bootstrap replications. The species targeted in this study were highlighted in a black box.

### Experimental Design, Materials and Methods Whole genome sequencing

Fresh sago palm leaves were harvested from the tree chosen by Lim et al. (2020a) for whole chloroplast sequencing in which this tree has been previously verified by our plant expert team from the living germplasm of Universiti Malaysia Sarawak East Campus (1°27′52.3″N, 110°27′04.7″E). The Illumina reads of sago palm were obtained from Lim et al. (2021) which was sequenced using Illumina HiSeq X.

In order to produce Oxford Nanopore long reads, the genomic DNA was first extracted from fresh sago palm leaves using high-salt SDS approach. About 5 μg of unfragmented genomic DNA libraries was prepared using LSK110 kit according to manufacturer’s protocols. The resultant libraries were sequenced on Oxford Nanopore PromethION platform. Reads were basecalled employing high-accuracy mode in Guppy version 4 and only passed reads were uploaded to the NCBI SRA database. The minimum cutoff length was set at 500 bp.

### Hybrid genome assembly and genome annotation

The Maryland Super-Read Celera Assembler v.3.2.2 was utilized to perform the *de novo* hybrid genome assembly of the sago palm. The sago palm hybrid genome assembly was conducted emulating that of Tan et al. (2018).

The prediction of all protein-coding genes was conducted using Augustus. A BUSCO analysis was performed on the annotated sago palm genome dataset utilizing the embryophyta-specific dataset as reference. TransDecoder v5.5.0 was employed to extract the protein coding sequences prior to the clustering at 98% similarity of protein using cdhit v4.7 (-g 1 -c 98). The functional annotation was done using eggNOGmappr (evolutionary genealogy of genes: Non-supervised Orthologous Groups) with a 0.001 minimum E-value.

### Phylogenetic analysis

All available full monocot genomes (with complete records of coding sequences and proteins) found within the GenBank public database were downloaded, totaling to 20 monocots and two dicot outgroups (only one representative was selected for each duplicated species). BUSCO analysis was conducted to all full genomes downloaded to extract all single-copy proteins. These proteins were aligned using Muscle prior to the maximum likelihood phylogenetic tree plotting using iqtree with 1000 bootstrap replications. The tree was visually improved using FigTree software.

## CRediT author statement

**Lim Whye Kit Leonard:** Data Curation, Writing-Original Draft. **Melinda Lau Mei Lin:** Data Curation. **Chung Hung Hui:** Conceptualization, Funding acquisition, Writing-Review and Editing. **Hasnain Hussain:** Writing-Review and Editing. **Gan Han Ming:** Methodology, Conceptualization, Writing-Review and Editing.

## Acknowledgments

This work was fully funded by Tun Openg Chair Research Grant F07/TOC/2068/2021 awarded to H. H. Chung by Centre for Sago Research (CoSAR), UNIMAS.

## References

Abbas, B., Dailami, M., Santoso, B., & Munarti. (2017). Genetic variation of sago palm (Metroxylon sagu Rottb.) progenies with natural pollination by using RAPD markers. Natural Science, 9(4), 104–109.

Austin, C. M., Tan, M. H., Harrisson, K., Lee, Y. P., Croft, L. J., Sunnucks, P., Pavlova, A., & Gan, H. M. (2017). De novo genome assembly and annotation of Australia’s largest freshwater fish, the Murray cod (Maccullochella peelii), from Illumina and Nanopore sequencing read. Gigascience, 6(8), 1–6.

Karim, A. A., Tie, A. P. L., Mana, D. M. A., & Zaidul, I. S. M. (2008). Starch from the sago (Metroxylon sagu) palm tree: Properties, prospects, and challenges as a new industrial source for food and other uses. Comprehensive Reviews in Food Science and Food Safety, 7(3), 215–228.

Lim, L.W.K., & Chung, H.H. (2020). Salt tolerance research in sago palm (Metroxylon sagu Rottb.): Past, present and future perspectives. Pertan. J. Trop. Agric. Sci., 43 (2), 91–105.

Lim, L.W.K., Chung, H.H., & Hussain, H. (2020a). Complete chloroplast genome sequencing of sago palm (Metroxylon sagu Rottb.): Molecular structures, comparative analysis and evolutionary significance. Gene Rep., 19, 100662.

Lim, L.W.K., Chung, H.H., & Hussain, H. (2020b). Organellar genome copy number variations and integrity across different organs, growth stages, phenotypes and main localities of sago palm (Metroxylon sagu Rottboll) in Sarawak, Malaysia. Gene Reports, 21, 100808.

Lim, L.W.K., Chung, H.H., Hussain, H., & Bujang, K. (2019). Sago palm (Metroxylon sagu Rottb.): now and beyond. Pertan. J. Trop. Agric. Sci., 42 (2), 435– 451.

Lim, L.W.K., Chung, H.H., Hussain, H., & Gan, H.M. (2021). Genome survey of sago palm (Metroxylon sagu Rottboll). Plant Gene, 28, 100341.

Nisar, M., & Hussain, H. (2020). The protein extraction method of Metroxylon sagu leaf for high-resolution two-dimensional gel electrophoresis and comparative proteomics. Chem. Biol. Technol. Agric,. 7, 14.

Tan, M. H., Austin, C. M., Hammer, M. P., Lee, Y. P., Croft, L. J., & Gan, H. M. (2018). Finding Nemo: hybrid assembly with Oxford Nanopore and Illumina reads greatly improves the clownfish (Amphiprion ocellaris) genome assembly. Gigascience, 7, 1–6.

Wee, C. C., & Roslan, H. A. (2012). Expressed sequence tags (ESTs) from young leaves of Metroxylon sagu. 3 Biotech, 2(3), 211–218.

Yoneta, R., Okazaki, M., & Yano, Y. (2006). Response of sago palm (Metroxylon sagu Rottb.) to NaCl stress. Sago Palm, 14(1), 10–19.

